# Pumping Venom: Valvilli Architecture and Implications for Stinger Functionality

**DOI:** 10.1101/2025.08.28.672817

**Authors:** Alexandre Casadei-Ferreira, Johan Billen, Sebastian Büsse, Julian Katzke, Riou Mizuno, Adrian Richter, Evan Economo

## Abstract

The internal structure of the insect stinger plays a central role in venom delivery, yet critical components in this process remain poorly understood. Of these, the valvilli, a pair of articulated structures within the valve chamber that act as ‘flaps’, have often been overlooked or described only superficially. Here, we use high-resolution micro-computed tomography (micro-CT), histological sectioning, and confocal laser scanning microscopy (CLSM) to characterize the fine morphology and material composition of the cuticle of ant valvilli. We report structural differentiation within the valvilli, including sharply delimited material zones that may correspond to distinct mechanical properties. CLSM imaging highlights variation in sclerotization, resilin distribution, and ultrastructure, indicating that the valvilli are not uniform flap-like elements but rather complex, partially deformable structures. Furthermore, comparative micro-CT scans of stingers fixed in different actuation states indicate the involvement of a non-antiphasic pattern of valvilli movement during stinger deployment, challenging the current understanding of valvilli-assisted pumping based on rhythmic, alternating movements, as inferred from European honeybees. These findings offer new anatomical insight into the architecture of the ant stinger and provide a refined morphological basis for future studies of venom delivery mechanisms in Hymenoptera; to also foster our understanding of the complex functionality of micro-scale injection and pumping systems in insects.

## 1. Introduction

Venom injection in Hymenoptera encompasses a diversity of morphological solutions, each evolved to achieve the critical biological goal of delivering venom effectively. Despite the apparent simplicity of venom injection through a sharpened pointy structure (Morse and Snodgrass, 2018; Snodgrass, 1983, 1956, 1933), close examination reveals substantial variation in the underlying biomechanical strategies among different hymenopteran lineages. While previous research has investigated various aspects of external stinger morphology (e.g., Hermann and Chao, 1983; Kugler, 1979; Kumpanenko et al., 2022; Kumpanenko and Gladun, 2024, 2018; Lieberman et al., 2022; Ramirez-Esquivel and Ravi, 2023; Snodgrass, 1956, 1933; Stetsun et al., 2019; Stetsun and Matushkina, 2020) and the biochemical composition of venom (e.g., Touchard et al., 2016, 2015), the mechanisms of venom delivery remain less understood. Particularly in taxa where differentiated internal structures modulate venom pumping, structural and functional relationships are unclear.

Within Aculeata (ants, bees, and wasps), two fundamental mechanisms for venom injection have been described: “pumping” and “injection” (Stetsun et al., 2019; Stetsun and Matushkina, 2020; Van Marle and Piek, 1986). The pumping mechanism, which is characteristic of bees (Apoidea) and ants (Formicidae), involves specialized internal structures known as valvilli (Lieberman et al., 2022; Snodgrass, 1933). These flap-like valvilli, positioned on the lancets (Lieberman et al., 2022), move rhythmically during venom delivery, actively generating hydraulic pressure to drive venom posteriorly through the venom canal (Ramirez-Esquivel and Ravi, 2023; Snodgrass, 1933). Notably, despite the widespread use of the term “valves” or “valvilli” to describe these internal structures, a recent study (Ramirez-Esquivel and Ravi, 2023) argues that their functional role in venom injection is more similar to that of collapsible pistons than true valves. Ramirez-Esquivel and Ravi (2023) hypothesized that valves regulate directionality without generating force, whereas pistons actively create positive pressure to propel fluid. Consequently, the term “valve” here arises from morphological homology rather than strict functional analogy, introducing potential confusion when discussing the biomechanics of venom injection. However, for consistency with existing literature, we continue to use the term “valvilli” throughout this manuscript (Lieberman et al., 2022; Snodgrass, 1956).

In contrast to the pumping mechanism of ants and bees, most wasps (i.e., Vespidae, Pompilidae, Crabronidae) employ an injection-type mechanism that lacks valvilli (Kumpanenko and Gladun, 2024, 2018; Stetsun et al., 2019; Stetsun and Matushkina, 2020; Van Marle and Piek, 1986). In these groups, the venom duct opens in the posterior region of the bulb and venom is propelled entirely through musculature-driven hydraulic pressure around the venom reservoir and duct (Van Marle and Piek, 1986). Interestingly, several wasp families, such as Mutillidae (velvet ants) (Kumpanenko et al., 2022), possess valvilli-like structures (Gadallah and Assery, 2004; Kumpanenko et al., 2022) resembling those found in bees and ants, although the extent to which these contribute to venom pumping remains unclear. This variation suggests that the presence of valvilli and the use of pumping versus injection mechanisms do not follow a simple phylogenetic pattern but instead represent diverse evolutionary solutions to venom delivery.

While the “pumping” and “injection” mechanisms are often presented as distinct and mutually exclusive systems, Robertson (1968) noted that in some bees, a small but distinct muscle remains associated with the venom gland. Although this muscle is far less developed than the strong gland-encasing musculature found in injection-type wasps, its presence suggests that venom ejection in these taxa may not rely solely on pressure driven by the valvilli. This indicates the possibility of intermediate or mixed systems within Aculeata, where valvilli motion and residual glandular musculature contribute to venom injection. Such arrangements could represent retained ancestral traits or transitional stages in the evolutionary shift between injection-type and pumping-type stingers, potentially with individual adaptive benefits and trade-offs.

In addition to the functional diversity seen in venom injection strategies, the morphological characteristics of valvilli themselves exhibit considerable variation across Hymenoptera (Quicke et al., 1992). While valvilli are generally articulated chitinous flaps located on the lower ovipositor or stinger valves, wide variability in their number, position, and structural complexity is documented (Quicke et al., 1992). Detailed surveys using light and electron microscopy have identified valvilli in most Ichneumonoidea and various aculeate lineages, while they are absent from other groups, including Symphyta, Evanioidea, and Cynipoidea (Quicke et al., 1992). Among Aculeata, valvilli typically appear as complex, paired structures mounted on a distinct base, an arrangement structurally adapted to facilitate venom pumping (Kumpanenko et al., 2022; Ramirez-Esquivel and Ravi, 2023; Snodgrass, 1956). In parasitoid Ichneumonoidea, by contrast, valvilli are structurally simpler and lack a defined base, likely reflecting their primary function in egg laying rather than venom injection (Quicke et al., 1992). Phylogenetically, the distribution and structural variation of valvilli appear significant, especially within Ichneumonidae and Braconidae, where differences in valvillus morphology correlate closely with broader evolutionary relationships (Quicke et al., 1992). The morphological and functional diversity of valvilli raises intriguing questions about the evolutionary pressures that shaped these features and highlights valvilli as valuable traits for understanding hymenopteran phylogeny and life history. This emphasizes the need for detailed anatomical and functional studies of valvilli across different hymenopteran groups to provide context for exploring their biomechanical roles in venom injection and oviposition systems.

The variation in venom-pumping strategies and valve morphology raises compelling evolutionary and functional questions. Specifically, if muscle-based venom ejection is energetically or mechanically efficient, what evolutionary pressures favored the development of valvilli in ants, bees, and some wasps? Conversely, if the valvilli-driven mechanism represents an optimization of some parameter of venom injection (e.g., dosage control, resistance to backflow, or sustained performance over repeated use), why have many wasp lineages developed the muscular-driven system? One possibility is that muscle-driven ejection offers faster, high-volume discharge, well-suited to rapid defense or prey subjugation, whereas valvilli pumping enables finer control, consistency, or pressure regulation across multiple stings. Answering these questions requires detailed comparative analyses of internal stinger morphology, material properties of the cuticle, and an understanding of the underlying fluid dynamic principles operating at microscopic scales in the different systems.

To date, most functional inferences about valvilli have assumed a simple “flap-like” structure capable of producing alternating movement to generate pressure for venom injection. However, no study has yet examined these structures using a combination of modern imaging techniques capable of revealing fine-scale morphology, material differentiation, and dynamic configuration. Here, we discuss in detail the internal stinger morphology of ants, using high-resolution micro-CT scanning, histological sectioning, and confocal microscopy (CLSM). Our goal is to characterize the three-dimensional architecture and material composition of the valvilli in unprecedented detail. In doing so, we identify characteristics that deviate from the existing “flap-like” paradigm, including differentiated cuticular material zones and apparent interdigitation that may facilitate more complex motion.

Furthermore, our comparative micro-CT data from specimens fixed at alternating phases of the sting cycle suggest that valvilli do not operate in a simple antiphasic rhythm but may exhibit both synchronous and asynchronous movements across the contraction cycle. These findings warrant reconsideration of how valvilli operate during venom injection and highlight the importance of fine-scale structural analysis in interpreting functional anatomy. By refining our understanding of this specialised structure, this study contributes new insight into both the morphology and potential biomechanical roles of the valvilli in ant stingers in particular, and in Hymenoptera more generally.

## 2. Materials and Methods

### a. Specimens Acquisition

For comparative morphological analyses, we used freshly collected specimens of *Diacamma* nr. *indicum* (Formicidae: Ponerinae) from a population in Okinawa, Japan, to ensure consistency in body size and morphology. Workers (adults, body size 8–16 mm) were collected from Kenmin no Mori (Onna, Kunigami District, Okinawa; 26.5063ºN, 127.9090ºE) and maintained in laboratory nests for direct access to fresh material. To capture variation in stinger posture during venom injection, individuals were manually provoked to sting and subsequently frozen using nitrogen spray. The resulting specimens had the stinger in either an extended (contracted/activated) or resting (relaxed) position (Figure 1A). Two individuals showing clear, opposing stinger configurations were selected for high-resolution micro-computed tomography (micro-CT). Additional individuals from the same colony were used for confocal laser scanning microscopy (CLSM). For CLSM, fresh specimens were dissected to isolate the stylet from the stinging apparatus and preserved in 99% ethanol until imaging. Specimens for histological sectioning were collected separately by JB in Okinawa and were not part of the main laboratory colony (see the Histology Section below for preparation details). Comparative analysis of internal stinger morphology among workers is based exclusively on micro-CT datasets. Details of staining duration, scan settings, and reconstruction parameters are provided in the Micro-CT Scanning section.

**Figure 1.**
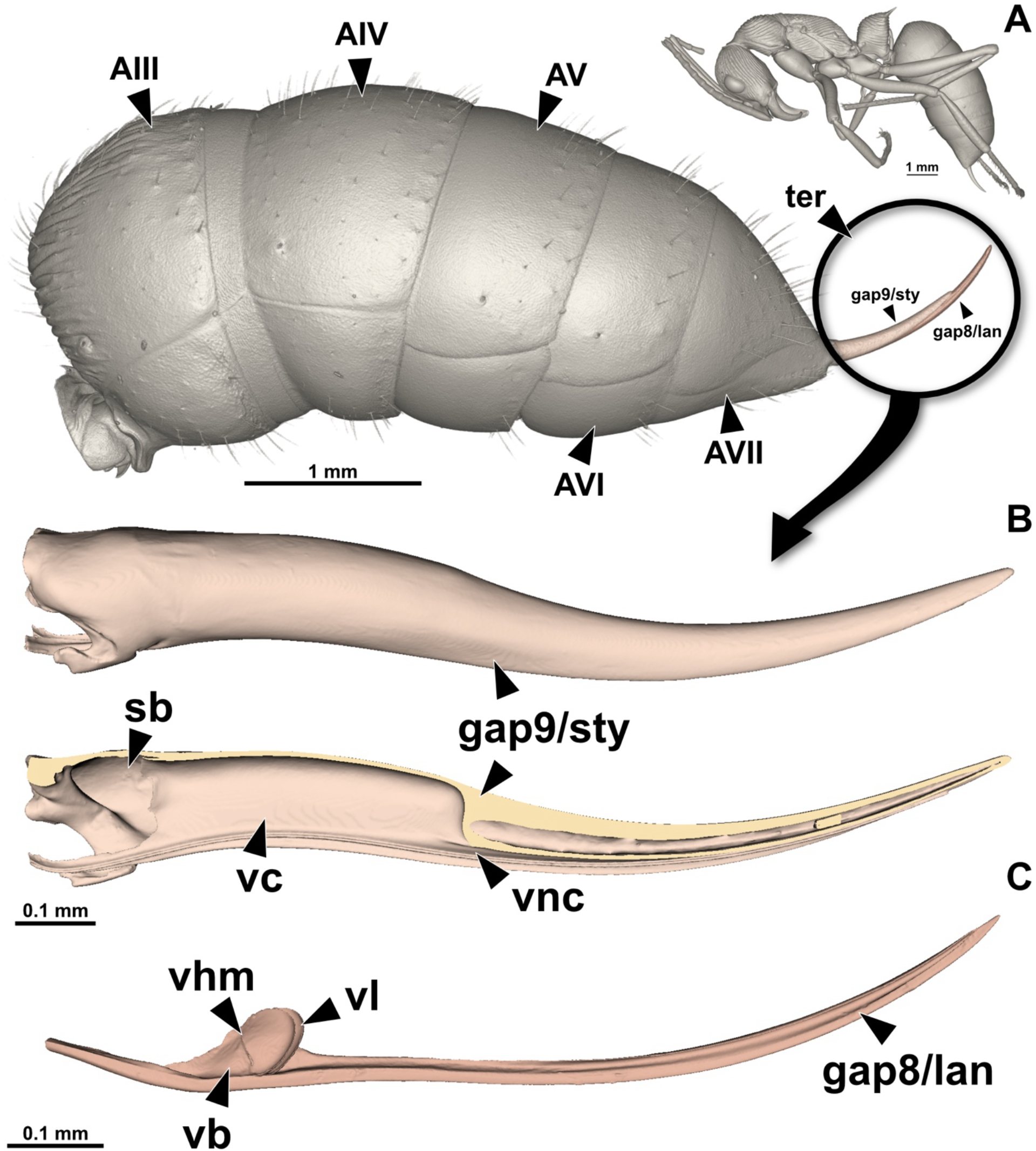
Three-dimensional reconstruction of the stinger apparatus in *Diacamma* nr. *indicum*. (A) External habitus showing abdominal segments AIII–AVII and the terebra (*ter*). The full-body 3D reconstruction was generated from micro-CT data accessed at the AntScan database (specimen ID: CASENT0745048; freely available at https://www.antscan.info). (B) 3D reconstruction of the stinger, including the stinger base (*sb*), stylet (*gap9/sty*, gonapophyses IX), and valve chamber (*vc*). (C) Close-up of a single lancet (*gap8/lan*, gonapophyses VIII) with associated internal structures: valvillus base (*vb*), hinge membrane (*vhm*), valvillus lobe (*vl*), and the venom channel (*vnc*).

### b. Histological Sections

For histological examination of the stinger, the metasoma of *D*. nr. *indicum* workers was carefully separated by transverse dissection between the 3^rd^ (AIII) and 4^th^ (AIV) abdominal segments. This approach facilitated effective penetration of fixatives and embedding media. Tissues were immediately fixed in cold 2% glutaraldehyde, buffered with 50 mM sodium cacodylate and supplemented with 150 mM saccharose to preserve structural integrity. Postfixation was done using 2% osmium tetroxide prepared in the same buffer. Following fixation, specimens were dehydrated through a graded acetone series and embedded in Araldite® epoxy resin. Serial semithin sections (1 μm thickness) were prepared using a Leica EM UC6 ultramicrotome (Leica Microsystems, Wetzlar, Germany) at the Zoological Institute, KU Leuven, Belgium. Sections were stained with a 0.1% methylene blue and thionin solution to enhance contrast for light microscopy and structural identification.

### c. Confocal Laser Scanning Microscopy (CLSM)

To assess the material composition of the valvilli and adjacent regions of the stinger in *D*. nr. *indicum*, we employed confocal laser scanning microscopy (CLSM) using a Leica Stellaris 8 system (Leica Microsystems, Wetzlar, Germany) at the Zoological Collection of the University of Rostock, Germany. This technique leverages the autofluorescent properties of cuticular components to infer relative differences in mechanical and structural characteristics based on cuticular material composition, such as sclerotization and resilin content (after Michels and Gorb, 2012). While this method allows qualitative mapping of relative material composition within the stinger, particularly the valvilli, it does not yield absolute quantitative values. Interpretations of cuticle properties are therefore limited to relative fluorescence intensity and spatial distribution (e.g. Josten et al., 2022). Although CLSM has previously been used to image stinger structures (Ma et al., 2023; Stetsun et al., 2019; Stetsun and Matushkina, 2020), this study represents, to our knowledge, the first application of the method targeting the valvilli.

Dissected stingers were mounted using the “sandwich” slide technique (Barbosa et al., 2014), in which the specimen is placed between two coverslips with play-dough spacers to prevent deformation. A drop of pure glycerin was used as a mounting medium to preserve tissue flexibility during imaging. Autofluorescent signals were acquired using three laser excitation wavelengths selected to target different material classes within the cuticle. Sclerotized cuticle was excited at 633 nm, corresponding to rigid, highly cross-linked chitin-protein complexes. Flexible cuticle was excited at 488 nm, representing areas with intermediate mechanical properties. Resilin-rich regions, associated with high elasticity, were excited at 405 nm (cf. Fig. 1 in Büsse and Gorb, 2018). Emitted light was detected in separate channels as grayscale images based on photon counts.

To maximize visual contrast and facilitate interpretation, we assigned each excitation channel a distinct false color: cyan (405 nm, resilin dominated), yellow (488 nm, flexible cuticle), and magenta (633 nm, sclerotized cuticle). This CMY palette offers high visual discrimination while avoiding overlap or saturation, providing a more aesthetically coherent and colorblind-friendly alternative to the conventional RGB-based schemes. Composite images were generated using ImageJ Fiji v1.54 software (National Institutes of Health, USA).

### d. Micro-CT Scanning

High-resolution X-ray micro tomography was performed using a Zeiss Xradia 510 Versa 3D X-ray microscope operated with the Zeiss Scout-and-Scan Control System at the Imaging Section, Okinawa Institute of Science and Technology Graduate University, Japan. Before scanning, all specimens were stained in a 4% elemental iodine (I_2_) solution dissolved in 99% ethanol for 48 hours to enhance soft tissue contrast. Following staining, samples were rinsed in clean 99% ethanol for at least 12 hours to remove excess iodine and standardize contrast conditions.

Scanning parameters were optimized for high absorption contrast and satisfactory resolution of internal structures. A low voltage setting of 40 kV was used to maximize contrast in iodine-stained soft tissues. To improve edge definition and visualization of subtle structural differences, the source–detector distance was slightly increased to enhance phase contrast capabilities inherent to the system. Each scan comprised 2001 X-ray projections acquired over a full 360° rotation using 4x and 20x magnification. Exposure times were adjusted individually to produce approximately 4000 intensity counts in the most highly absorbing tissues, ensuring consistent brightness and signal across samples.

Scans were reconstructed using Zeiss software and exported for post-processing. The reconstructed volumes were converted into NRRD format (“*nearly raw raster data*”) and imported into 3D Slicer v5.8.1 (Brigham and Women’s Hospital, USA) (Fedorov et al., 2012). 3D renders were generated using the *Colorize Volume* module from the 3D Slicer Sandbox extension for figure preparation.

## 3. Results

In *Diacamma* nr. *indicum*, the ninth gonapophyses are medially fused and form the distal part of the stinger base, including the bulb, the valve chamber, followed by the stylet portion of the terebra (Figure 1B). The bulb and valve chamber occupy the proximal region to the terebra, enclosing the space in which the venom-pumping structures are located. Within this chamber are the opposing valvilli, each projecting medially from one of the paired lancets (eighth gonapophyses). In gross view, each valvillus can be divided into three recognizable parts. At its origin on the lancet is a stout basal section (here referred to as the valvillus base, *vb*; Figure 2), which is internally hollow and provides the main attachment to the cuticular wall of the lancet. Two elongated lobes extend distally from the base (valvillus lobes, *vl*; Figure 2), which project into the valve chamber toward the midline. The connection between the base and the lobes is formed by a flexible cuticular zone (valvillus hinge membrane, *vhm*; Figure 2) that allows the lobes to pivot or flex relative to the base. In situ, the paired valvilli face each other across the central lumen of the valve chamber, occupying much of its dorsal space, with their lobes aligned so that they can approach or slide past one another during stinger actuation. This arrangement positions the valvilli ideally for modulating the transfer of venom from the bulb into the venom canal and finally injecting it into the target.

**Figure 2.**
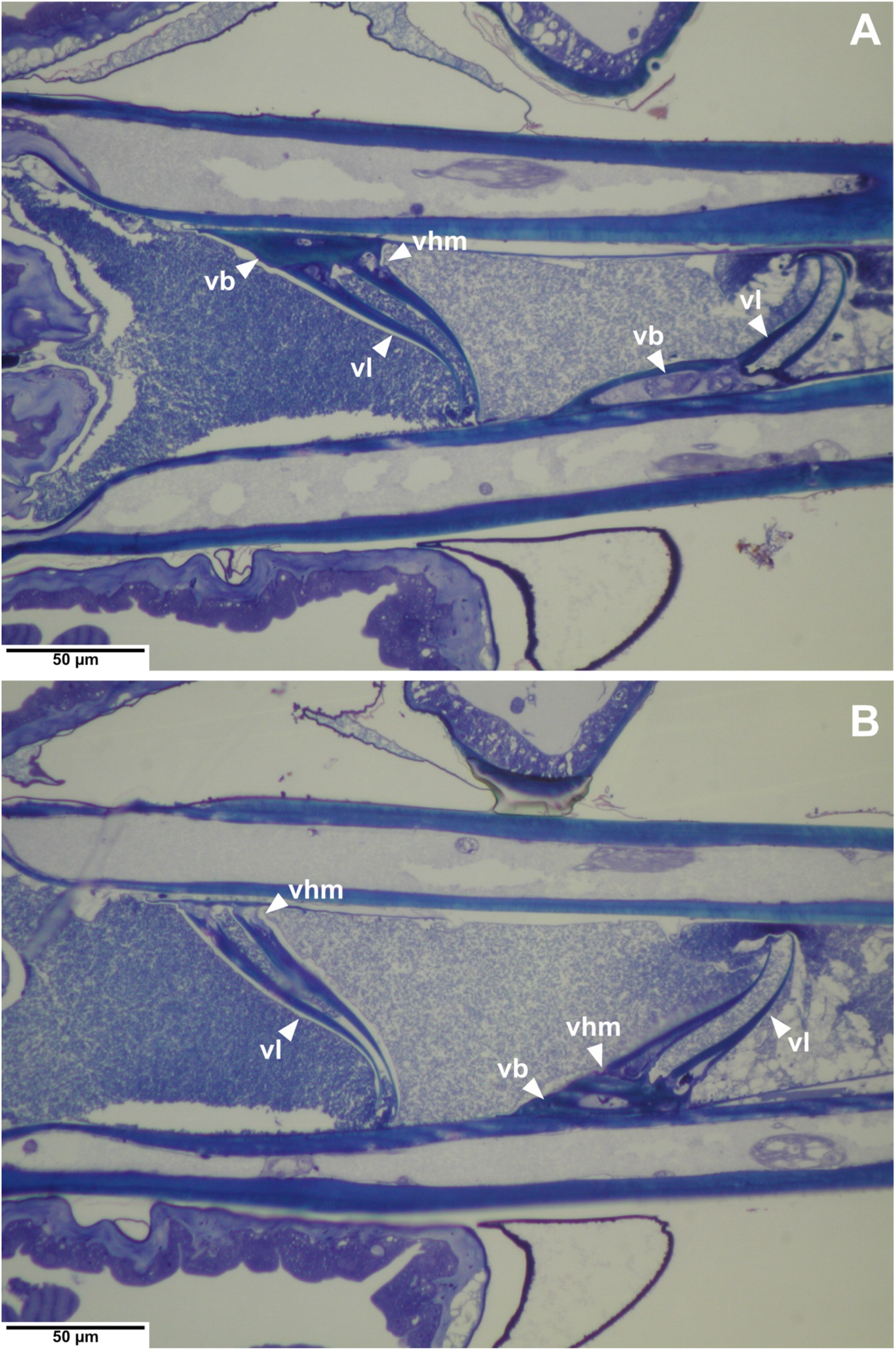
Coronal histological sections of the stinger showing valvillus organization. (A, B) Histological sections through the stinger base of *Diacamma* nr. *indicum*, viewed in coronal (dorsoventral) orientation. Sections capture different dorsoventral levels: (A) more ventral; (B) more dorsal. Arrows indicate key components of the valvillus: valvillus base (*vb*), valvillus hinge membrane (*vhm*), and valvillus lobe (*vl*).

Confocal laser scanning microscopy (CLSM) reveals distinct differences in the autofluorescence patterns of the stinger components, allowing qualitative assessment of the material composition of their cuticle. The region of the lancet bearing the valvillus originates within a strongly sclerotized portion of the stylet wall (magenta signal; Figure 3), which extends into the basal section of the valvillus itself. The valvillus base (*vb*) appears predominantly magenta (Figure 3), indicating a stiff, sclerotized cuticle, consistent with its structural role in anchoring the element to the lancet wall. In contrast, the valvillus hinge membrane (*vhm*) shows a mixed “green” signal (Figure 3), interpreted as an overlap of cyan and yellow channels, indicating a flexible chitin– resilin composite that is likely more deformable than stiff sclerotized cuticle, but less elastically responsive than resilin-rich regions. This zone appears to act as the principal pivoting point for lobe movement. The valvillus lobes (*vl*) are largely magenta (Figure 3), indicating a rigid composition, but CLSM revealed a distinct internal organization of the material composition within the lobes with fine, longitudinally orientated regions extending distally from the hinge membrane toward the apical margin (Figure 3). At higher magnification, the distal margin of the *vl* presents a segmented or “eyelash-like” appearance, with rigid magenta elements embedded within a narrow band of mixed-signal flexible cuticle (Figure 3B). The longitudinally oriented regions appear continuous with these marginal elements, producing a transition from the predominantly rigid central surface (core) of the lobe to a narrower flexible zone at its edge (Figure 3B). The two regions are differentiated by both color signal and structural organization.

**Figure 3.**
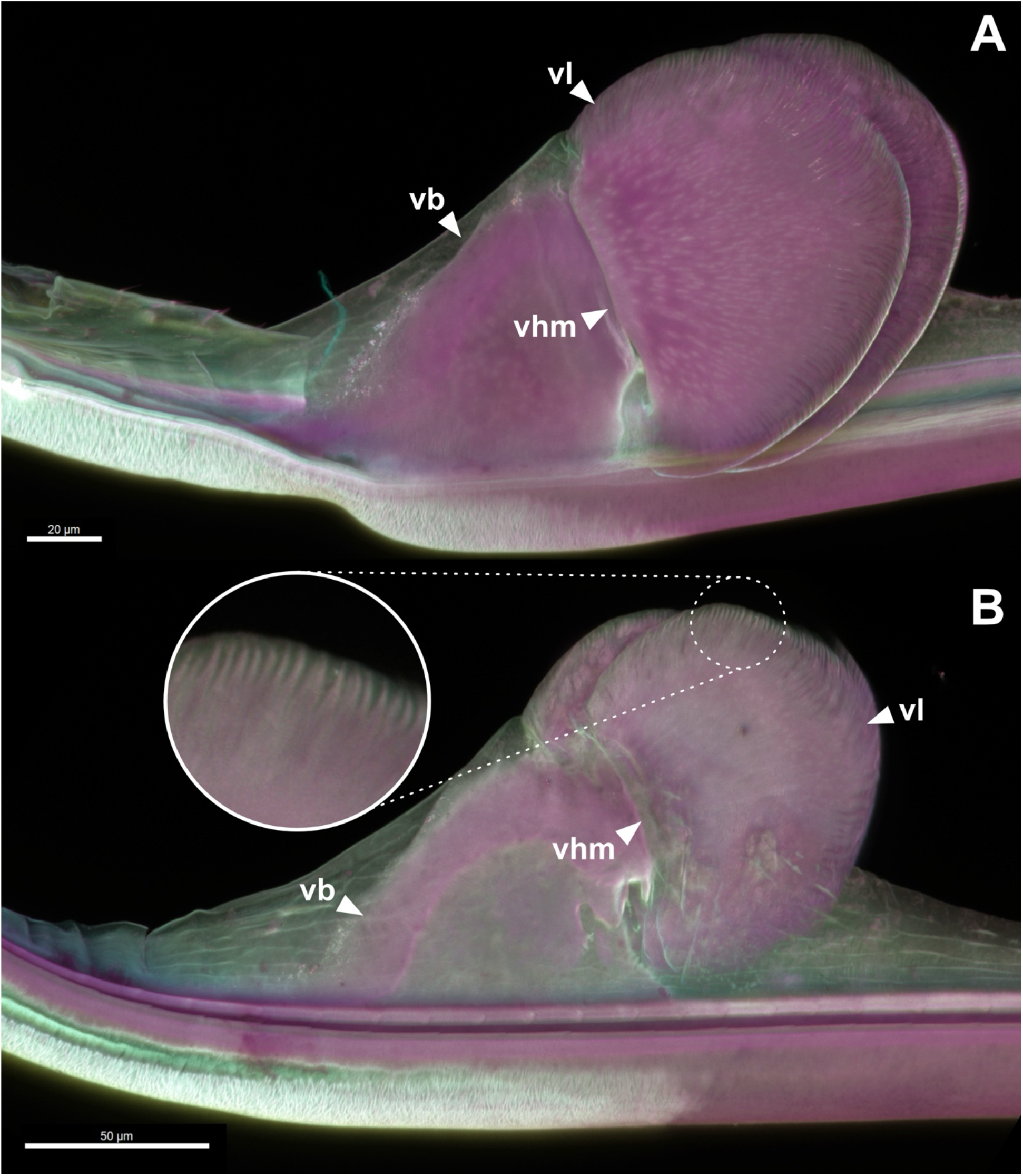
Material composition and internal structure of the valvilli revealed by confocal laser scanning microscopy. (A, B) CLSM maximum intensity projections of the valvillus-bearing region of the lancet in *Diacamma* nr. *indicum*, showing autofluorescent material signals assigned to sclerotized cuticle (magenta, 633 nm), flexible cuticle (yellow, 488 nm), and resilin-dominated regions (cyan, 405 nm). (A) Median view showing the valvillus base (*vb*), hinge membrane (*vhm*), and lobe (*vl*). The *vb* exhibits a strong magenta signal, indicating a rigid, sclerotized cuticle, while the *vhm* shows a mixed green signal (cyan + yellow), suggesting a flexible chitin–resilin composite. The lobe (*vl*) is predominantly rigid but contains fine, longitudinally oriented internal bands of magenta signal extending distally from the hinge zone. (B) Lateral view with higher magnification of the distal lobe margin, revealing a segmented or “eyelash-like” edge where rigid elements (magenta) are embedded within a narrow band of flexible cuticle (green signal).

Micro-CT scans of the stingers fixed in different actuation states reveal marked variation in the relative position and orientation of the paired valvilli within the valve chamber (Figure 4C). In most reconstructions, the *vhm* was not clearly resolved and frequently appeared as a gap between the *vb* and *vl* (Figure 4C). The proximal bulb is consistently broader than the valve chamber (*vc*), which tapers distally toward the venom canal (Figure 4B and C). In several specimens, the narrowing corresponds with a posture in which the valvilli are pressed closely together within the chamber, their lobes deflect laterally toward the chamber wall due to the flexible *vhm*, which forms the articulation between the base and the lobes (Figure 4B2 and C2). In this configuration, both valvilli might act together, as our micro-CT images recover them displaced distally (Figure 4C3), compressed against the distal wall of the chamber. Histological sections used in this study show another different posture (Figure 2), with the valvilli offset, one displaced proximally and the other remaining distally, creating a visible gap between their medial margins. These two observed configurations (i.e., synchronous and asynchronous) correspond to the positional stages illustrated in the movement sequence diagram (Figure 4A and B; Supplementary Movie 1).

**Figure 4.**
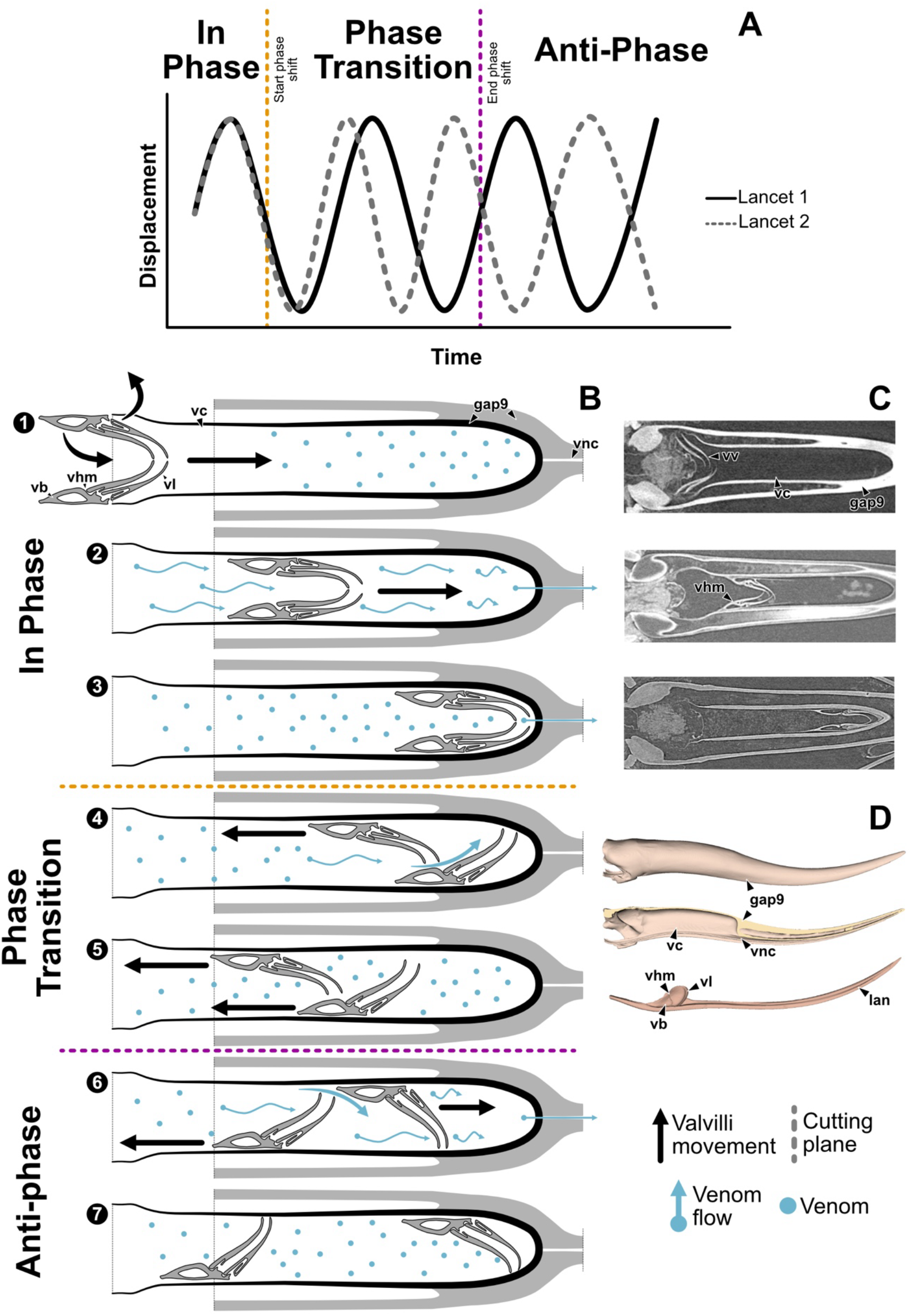
Proposed model of the stinging dual-phase motion cycle in *Diacamma* nr. *indicum*. (A) Schematic representation of hypothesized phase transitions during a stinging cycle. The cycle begins with both lancets (and attached valvilli) moving distally in-phase, compressing against the valve chamber. During the return stroke, the motion gradually shifts to an antiphasic pattern, enabling rhythmic valvillus-based pumping of venom. (B) Illustration of valvilli mechanics in coronal section. When displaced in-phase (1–3), the rigid lobes are deflected laterally and press against the chamber wall, forming a seal. In the transition stage (4–5), one valvillus retracts while the other remains advanced, allowing venom to refill the chamber and initiating the anti-phase stage (6–7). (C) Coronal micro-CT slices of stingers fixed at different actuation states, all showing in-phase configurations with paired valvilli (*vv*) displaced distally within the narrowed valve chamber. The hinge membrane (*vhm*) is not resolved but inferred based on the gap between the valvillus base (*vb*) and lobe (*vl*). Note variation in lobe orientation and chamber tapering along the distal-proximal axis. (D) 3D reconstruction of the stinger, showing the stinger base (*sb*), stylet (*gap9/sty*, gonapophyses IX), and valve chamber (*vc*), and a detail view of a single lancet (*gap8/lan*, gonapophyses VIII) with associated valvillus structures, including base (*vb*), hinge membrane (*vhm*), lobe (*vl*), and venom channel (*vnc*). Collectively, the observed postures correspond to the two functional stages of the proposed dual-phase pumping mechanism.

## 4. Discussion

The internal pumping mechanism of the hymenopteran stinger has long been interpreted within a relatively narrow morphological framework. Early foundational works (Snodgrass, 1933; Van Marle and Piek, 1986) and more recent analyses (Ramirez-Esquivel and Ravi, 2023) describe the valvilli as flap-like cuticular projections that operate in strict antiphase motion to drive venom from the bulb into the venom canal. This view, derived largely from observations in the European honeybee, has been broadly applied to other Aculeata without equivalent fine-scale anatomical data. In particular, the structural complexity of the valvilli, and especially the organization of their lobes, has remained undescribed, and no study has directly documented their configuration across different sting postures in a single species. Our combined micro-CT, histological, and CLSM observations of *Diacamma* nr. *indicum* reveals architectural and compositional variation not captured in previous models.

The cuticle of the valvillus base, hinge membrane, and lobes each display distinct material zoning (Figure 2–4). Notably, the lobes are predominantly rigid but exhibit a distinctive internal arrangement of material zones (Figure 3). The hinge area itself shows a signal consistent with a flexible, possibly resilin-enriched zone. This arrangement resembles a syndesis (i.e., a flexible, resilin-rich connection between more rigid cuticular elements; see Josten et al., 2022) and may act as a pivot that permits repeated deflection of the rigid lobes on the valvillus base during actuation. By contrast, the distal arrangement of the lobes terminates in a comb-like edge, where rigid elements are embedded within a narrow, flexible band (Figure 3B). This arrangement and material transition suggest that the lobe margins may undergo localized deformation without a true joint, enabling them to seal against the chamber wall or slide past the opposing valvillus (Figure 4B and C). Moreover, our observations reveal a previously undocumented synchronous configuration in *Diacamma*, indicating that its functional cycle may combine both synchronous and alternating phases. This represents a broader kinematic repertoire than the canonical antiphasic model, underscoring the need to reassess pumping mechanisms across taxa and revisit long-standing assumptions about stinger biomechanics in Hymenoptera.

In more detail, histological sections (Figure 2) and micro-CT imaging (Figure 4C) of stingers fixed in alternate postures suggest two discrete configurations: an offset arrangement consistent with antiphasic motion and an in-phase stage. The first matches the expected offset arrangement of the valvilli (Figure 2), consistent with antiphasic motion (e.g., Ramirez-Esquivel and Ravi, 2023; Snodgrass, 1933). The second, not previously reported in Aculeata with pumping mechanisms, is an in-phase stage in which both lancets are displaced distally and their attached valvilli compressed within the narrowed distal valve chamber (Figure 4C3).

In this in-phase posture of the lancets, the hinge membrane acts as a pivot, permitting the rigid lobes to deflect toward the chamber wall, bringing their medial margins into close contact and forming a seal (Figure 4C2 and C3, Supplementary Movie 1). The combination of stiff core and flexible margin likely enhances this sealing effect by enabling precise conformity to the valve chamber. Such a configuration supports two complementary mechanical actions. First, during the in-phase stage (Figure 4A and C1–3, Supplementary Movie 1), the close apposition of the lobes within the narrowed distal valve chamber may produce an effective seal, maximizing pressure in the enclosed volume. This could drive the expulsion of a large bolus of venom in a single “push” before the cycle transitions to an antiphasic phase (Figure 4A and C6–7, Supplementary Movie 1). Second, the flexible margins and embedded rigid elements (Figure 3B) may permit one lobe to deform and slide past the other during antiphasic motion without losing integrity, sustaining a continuous pumping cycle (functionality of phasic and anti-phasic movement, see below).

The presence of an in-phase stage also has potential implications for the puncture mechanics of the stinger. As Anderson (2018) noted, puncture into ductile materials such as insect cuticle or vertebrate skin requires both fracture initiation and fracture propagation, and delivering sufficient energy to initiate the fracture is often the limiting factor, particularly for small-mass organisms. In *Diacamma*, the in-phase stage reflects synchronous movement of the paired lancets, so their sharpened tips advance together. This coordinated forward motion could transmit a concentrated mechanical load to the target, restricting deformation and facilitating fracture initiation (Anderson, 2018, 2005; Atkins, 2009; Blackman et al., 2013; Callister, 2004). Once the fracture is formed, the system could transition to an antiphasic mode, in which alternating lancet movements propagate the puncture while sustaining venom delivery.

Comparable alternating movements are well known in the ovipositors of parasitic wasps, which are homologous to the stingers of Aculeata (Cerkvenik et al., 2017; Eggs et al., 2018). In these taxa, push–pull mechanics between the individual valves allow insertion into resistant substrates while avoiding excessive axial loading that could lead to buckling of the slender structure (Cerkvenik et al., 2017). Although the stylets of *Diacamma* are more robust than many ichneumonid ovipositors, an alternating anti-phase stroke following the in-phase penetration could similarly reduce continuous axial force on the apparatus, lowering the risk of structural stress while maintaining forward progress. In ovipositors, differential protraction of the valves can also induce subtle steering of the tip by creating asymmetric reaction forces at the substrate interface (Cerkvenik et al., 2017). While such steering is unlikely to be a primary function in soft tissue stinging, alternating motion in *Diacamma* may nonetheless stabilize the puncture trajectory or compensate for minor lateral deflections during insertion. The underlying musculoskeletal arrangements that drive alternating motion in parasitoid wasps — antagonistic muscle pairs acting through lever-like valvifers — are structurally conserved across Hymenoptera (Eggs et al., 2018) and likely underpin the rapid and coordinated transitions between synchronous and alternating phases (Figure 4A and C4–5, Supplementary Movie 1) that we hypothesized in *Diacamma*.

Within Aculeata, the sequence and coordination of stinger movements show notable variation, with both fully alternating and partly synchronous kinematics occurring. For the European honeybee, a strictly alternating sequence is described (Snodgrass, 1933), in which the lancets advance and retract in turn, each anchoring via its barbs while the other moves forward (see Ramirez-Esquivel and Ravi, 2023 for further discussion). This ratcheting action progressively draws the stinger deeper with minimal resistance. Kumpanenko and Gladun (2018) documented a similar general pattern of alternating strokes in *Cryptocheilus versicolor* (Scopoli) (Pompilidae), but noted that specific structural characteristics, such as the wide anal arc of tergum IX and its articulation with the first valvulae, which creates a rigid connection between its two halves, can limit independent movement. This results in phases where both lancets advance or retract together. In other aculeate groups, including Vespidae, Formicidae, and Apoidea, the anal arc is much narrower, which permits the halves of tergum IX to shift slightly against each other (Kumpanenko and Gladun, 2018; Lieberman et al., 2022; Snodgrass, 1956). This increased mobility enables the first valvulae to operate alternately in valve-pump type stingers. These morphological observations emphasize that stinger motion in Aculeata is not fixed to a single coordination pattern but instead reflects a continuum from fully alternating to partly synchronous sequences.

The dual-phase coordination observed in *Diacamma*, which combines distinct synchronous and alternating stages, introduces an additional level of complexity to this spectrum, emphasising the adaptability of hymenopteran stinger kinematics. This mechanical flexibility also underpins how the in-phase and anti-phase stages will contribute differently to venom delivery. Movement patterns are probably directly related to the dynamics of venom pumping and flow. While the in-phase stage likely plays a key role in fracture initiation and initial high-pressure delivery, this posture also completely seals the valve chamber, preventing refill. As we have not observed an opening between the valvilli in the in-phase stage, an anti-phase mode is essential to break the seal and consistently replenish the valve chamber with venom. The rearward displacement of one lancet would create an opening between the lobes (Figure 4B4, Supplementary Movie 1), allowing venom to flow from the gland into the chamber. Thus, alternating back-and-forth movements might enable successive partial refills, maintaining a consistent supply to the venom canal.

The motoric complexity of the dual-phase pumping cycle contrasts with the “injection” mechanism of many wasp lineages, where valvilli are absent and venom is expelled by direct muscular compression of the reservoir (Kumpanenko and Gladun, 2024, 2018; Stetsun et al., 2019; Stetsun and Matushkina, 2020; Van Marle and Piek, 1986). This system is mechanically simpler and can deliver venom rapidly in large volumes, but it sacrifices the fine metering and potential dual-mode operation of a valve pump. The retention of valvilli in ants and bees may be favored where controlled delivery is advantageous, for example, in delivering small amounts of venom to multiple targets, modulating venom output depending on context, or maintaining pressure over extended injections. By contrast, the injection mechanism may be optimized in contexts where speed and volume outweigh precision, such as rapid defense or the subjugation of large prey. The fact that some wasp families (e.g., Mutillidae; Kumpanenko and Gladun, 2024) have retained valvilli suggests that selective pressures vary even within wasps, perhaps linked to prey handling, nesting biology, or the balance between defensive and predatory functions of the stinger. Conversely, the loss of valvilli in other lineages may reflect a shift towards maximizing delivery rate over control, reduced reliance on venom economy, or morphological constraints that make the valve chamber less effective.

The physical properties of the venom cocktail itself likely influence the efficiency of this refill- and-seal cycle. Although the mechanical aspects of stinger actuation are relatively well understood, the physicochemical properties of ant venom and their potential implications for flow at the microscale are not well studied. Proteomic and peptidomic surveys (Touchard et al., 2016, 2015) have shown that ant venoms are complex cocktails comprising not only low-molecular-weight compounds and enzymes but also a broad range of peptides, from short linear cytolytic peptides to larger disulfide-rich neurotoxins and dimeric structures. Many of these highly structured peptides may occur at relatively high concentrations, potentially increasing viscosity compared with simple aqueous solutions. Enzymatic components such as phospholipases, proteases, and hyaluronidases, common in many ant venoms, can further alter local tissue environments, but their contribution to intra-stinger fluid mechanics is unknown. At present, no direct measurements of ant venom rheology exist, leaving its flow behavior inside the stinger largely speculative. However, studies on microneedle and microfluidic systems show that in channels narrower than ∼100 µm, viscous drag dominates and surface tension can significantly resist liquid entry, especially for solutions containing larger biomolecules (Ita, 2020; Martanto et al., 2005). Under such conditions, even moderate increases in viscosity can markedly slow passive flow, making active “opening” phases crucial for rapid chamber refill.

These considerations show that beyond the mechanical distinctions between pumping and injection systems, the effectiveness of the stinger is also shaped by the associated glandular complex and its products. Robertson (1968) emphasized that the venom apparatus comprises a set of integrated glands whose morphology and function vary across Hymenoptera. The main venom gland produces the toxic secretion and delivers it to the bulb region of the stinger via a venom duct. This process is often aided by a specialised filtering or narrowing structure that can regulate the flow of particles (Callahan et al., 1959). Closely associated is the Dufour’s gland, which opens into the stinger bulb and can function as a lubricant source, a chemical adjunct to venom, or — in many ants and social bees — a producer of pheromones involved in alarm signaling or trail marking (Morgan, 2009). In some taxa, notably certain species of ants, the release of Dufour’s secretion is neurally coordinated with venom ejection, suggesting an active role during stinging rather than passive by-flow. The arrangement of these gland openings in relation to the valve chamber means that their output is subject to the same mechanical pressures generated during lancet motion, linking glandular secretion directly to sting kinematics. In this light, sting mechanics should be viewed as part of an integrated system in which glandular output, flow control, and mechanical delivery are coupled.

While it is clear that variation in the function of the sting apparatus exists across aculeates, it remains possible that some of this variation is due to the lack of understanding of the full systems in various species. While European honeybee studies focus on the strict antiphasic cycles of the piston-like or collapsible valve-like valvilli (Ramirez-Esquivel and Ravi, 2023; Snodgrass, 1933; Van Marle and Piek, 1986), this model may be incomplete. Our considerations of the in-phase stage in *Diacamma* indicate that this may present advantages beyond the precision metering and continuous flow generated by the anti-phase stage. An in-phase stage may occur in bees, but this has been overlooked because prior studies did not analyze stingers in multiple postures or used methodologies that precluded fine-scale temporal resolution. Detecting such a phase would require targeted observation of stingers in different actuation states across a range of taxa. Whether the dual-phase cycle in *Diacamma* reflects a lineage-specific innovation or a shared capacity enabled by conserved morphological traits remains unresolved. Determining its distribution and evolutionary origin will require comparative studies of valvilli architecture and sting kinematics across a wider phylogenetic range.

Although the *Diacamma* valvilli conform to the general aculeate body plan of a robust base and lobes articulating via a flexible hinge (Ramirez-Esquivel and Ravi, 2023; Snodgrass, 1933; Van Marle and Piek, 1986), the internal lobe organization and potential dual-phase venom delivery discovered here were not described in any other hymenopteran. These findings raise important questions about the evolutionary history of the pumping mechanism, like: Did the in-phase stage arise within Formicidae, or is it an ancestral condition of the pumping apparatus that has gone unnoticed? Is the fine-scale compositional specialisation of the lobes a unique adaptation, or has it not been detected in other groups due to methodological limitations? By integrating structural, compositional, and positional data, our study provides the most detailed morphological account of ant valvilli to date. It introduces a revised functional model that incorporates both in-phase and anti-phase dynamics (Figure 4 and Supplementary Movie 1). The coupling of rigid and flexible elements within a single lobe, combined with dual-mode motion, suggests that the ant valvillus apparatus is adapted to meet both the mechanical stress during penetration and the physiological requirements of venom delivery. Extending high-resolution imaging and multi-posture sampling to a broader range of taxa will be essential to determine whether this versatile pumping system is unique to ants, convergently evolved in other valvillus-bearing lineages, or ancestral to Aculeata and subsequently lost in some groups.

## 5. Conclusions

Our findings redefine the functional paradigm of aculeate venom-pumping mechanisms, departing from the canonical bee model, by showing that the *Diacamma* nr. *indicum* stinger operates in a dual-phase cycle, alternating between an in-phase, high-pressure stage and an anti-phase, refill stage. High-resolution micro-CT imaging of stingers fixed in alternate postures provided direct morphological evidence of the in-phase configuration, with both valvilli displaced distally and compressed together within the valve chamber. This cycle is enabled by a previously undescribed internal organization of the valvillus lobes, in which rigid structural support is coupled with locally deformable margins, balancing the mechanical demands of lancet-driven penetration with the physiological requirements of sustained venom delivery. As the most detailed morphological account of ant valvilli to date, these results expand the known functional repertoire of the pumping mechanism and highlight the need for comparative, high-resolution analyses across Hymenoptera to uncover the evolutionary origins and ecological drivers of this versatility. Targeted studies integrating *in vivo* kinematics with fine-scale morphological and material analyses will be essential to determine whether this dual-phase cycle is unique to ants or represents an overlooked ancestral condition within Aculeata with pumping mechanisms.

## Acknowledgments

We thank Fiorella Ramirez-Esquivel, Karen Kohama, Lakshmipriya Swaminathan, and Kazumi Toda-Peters for their insightful discussions throughout the development of this paper. We thank Christian Wirkner and Stefan Richter at the Zoological Collection of the University of Rostock for giving access to the Leica Stellaris 8 CLSM system, Maximilian Stüwe for support during CLSM image acquisition, and An Vandoren for making the histological sections. We thank the Okinawa Institute of Science and Technology Graduate University (OIST) Imaging Section for providing access to the Zeiss Xradia micro-CT scanner and Shinya Komoto for general support. We acknowledge the KIT Light Source for providing instruments at their beamlines, and we would like to thank the Institute for Beam Physics and Technology (IBPT) for operating the storage ring, the Karlsruhe Research Accelerator (KARA), which made it possible to generate Antscan data.

